# Traffic-derived particulate matter and angiotensin-converting enzyme 2 expression in human airway epithelial cells

**DOI:** 10.1101/2020.05.15.097501

**Authors:** L Miyashita, G Foley, S Semple, J Grigg

## Abstract

**Background:** The mechanism for the association between traffic-derived particulate matter less than 10 microns (PM_10_) and cases of COVID-19 disease reported in epidemiological studies is unknown. To infect cells, the spike protein of SARS-CoV-2 interacts with angiotensin-converting enzyme 2 (ACE2) on host airway cells. Increased ACE2 expression in lower airway cells in active smokers, suggests a potential mechanism whereby PM_10_ increases vulnerability to COVID-19 disease.

**Objective:** To assess the effect of traffic-derived PM_10_ on human airway epithelial cell ACE2 expression *in vitro*.

**Methods:** PM_10_ was collected from Marylebone Road (London) using a kerbside impactor. A549 and human primary nasal epithelial cells were cultured with PM_10_ for 2 h, and ACE2 expression (median fluorescent intensity; MFI) assessed by flow cytometry. We included cigarette smoke extract as a putative positive control. Data were analysed by either Mann-Whitney test, or Kruskal-Wallis with Dunn’s multiple comparisons test.

**Results:** PM_10_ at 10 μg/mL, and 20 μg/mL increased ACE2 expression in A549 cells (P<0.05, 0.01 vs. medium control, respectively). Experiments using a single PM_10_ concentration (10 μg/mL), found increased ACE2 expression in both A549 cells (control vs. PM_10_, median (IQR) MFI; 470 (0.1 to 1114) vs 6217 (5071 to 8506), P<0.01), and in human primary epithelial cells (0 (0 to 591) vs. 4000 (2610 to 7853), P<0.05). Culture of A549 cells with 5% cigarette smoke extract increased ACE2 expression (n=4, 0 (0 to 28) vs. 9088 (7557 to 15831, P<0.05).

**Conclusion:** Traffic-related PM_10_ increases the expression of the receptor for SARS-CoV-2 in human respiratory epithelial cells.

## Background

There is emerging evidence for an association between air pollution and COVID-19 disease caused by the pathogenic SARS-coronavirus 2 (SARS-CoV-2). For example, Zhu *et al* (1) using data from China reported that a 10 μg/m^3^ increase in particulate matter (PM) with an aerodynamic diameter of less than 2.5 microns (PM_2.5_) over the previous 2 weeks was associated with a 2.2% (95%CI 1.02 to 3.4) increase in newly confirmed COVID-19 cases, and Fattorini and Regoli (2) reported that long-term air-quality data from 71 Italian provinces significantly correlated with COVID-19 cases. However, there remains an urgent need for mechanistic research to delineate the biological plausibility for a link between air pollution and COVID-19 disease (3).

In order to initiate infection, SARS-CoV-2 must first engage with angiotensin-converting enzyme 2 (ACE2) expressed on airway cells (4). Angiotensin-converting enzyme 2 is a membrane-associated aminopeptidase, and viral infection is facilitated by an interaction between the extracellular portion of ACE2 and the receptor binding domain of the SARS-CoV-2 spike (S) glycoprotein. Cryogenic electron microscopy experiments show that SARS-CoV-2 has ten times the affinity to ACE2 compared with SARS-CoV, the virus responsible for the SARS epidemic (5). After engaging with ACE2, cellular serine protease TMPRSS2 primes SARS-CoV-2 RNA genome for cell entry (4). Analysis of single cell RNA-sequence datasets show that 1.4% of human type II pneumocytes express ACE2, with 0.8% of type II cells co-expressing both ACE2 and TMPRSS2 (6). In the upper respiratory tract, ACE2 is expressed by nasal ciliated epithelial and goblet cells (7). Recent studies suggest that ACE2 expression may be upregulated by mediators. For example, culture of primary human airway cells with interferon alpha 2 *in vitro* increases ACE2 transcripts (6).

Since the effect of traffic-related PM on the expression of ACE2 in human airway cell populations is not known we sought, in this study, to assess ACE2 expression in human airway epithelial cells exposed to traffic-derived PM_10_ *in vitro*.

## Methods

### Particulate matter

Traffic-derived PM_10_ was collected as dry particles using a high-volume cyclone placed within 2 metres of Marylebone Road, London, UK (8). Marylebone Road is one of the most polluted roads in Europe, with diesel trucks dominating near-road traffic-derived PM_10_ emissions (9). In order to obtain milligram amounts of PM_10_, sampling was done between 6 to 8 h per day on 10 occasions between May and September 2019 (i.e. before the UK lockdown). PM_10_ samples were pooled and stored at room temperature in a sterile glass container. An aliquot of PM_10_ was diluted in Dulbeccos phosphate-buffered saline (DPBS) to a final concentration of 1 mg/mL and stored as a master stock at −20°C.

### Cigarette smoke extract

Cigarette smoke extract (CSE) was collected onto a cotton filter through a peristaltic pump (Jencons Scientific Ltd., East Grinstead, UK) at a fixed rate from two Malborough red cigarettes, as previously described (10). Cigarette smoke extract was extracted after vortexing in 2 mL Dulbecco’s DPBS and stored at −80°C as 100% master stock.

### Airway cells

The human alveolar type II epithelial cell line A549 was purchased from Sigma-Aldrich (Poole, UK) and maintained in Dulbecco’s Modified Eagle Medium (DMEM) supplemented with fetal bovine serum (FBS) and penicillin-streptomycin (Lonza, Basel, Switzerland). Passage number was less than 20. Human primary nasal epithelial cells were obtained from a non-smoking, non-vaping, healthy adult donor using a dental brush, maintained in airway epithelial cell growth medium (AECGM), with supplement kit (PromoCell^®^, Heidelberg, Germany) with Primocin (InvivoGen, San Diego, USA), and stored at passage 1 cryogenically in freezing media (AECGM: 10% FBS, 10% DMSO). A vial of human primary nasal epithelial cells was thawed and aliquoted into multiple T25 cell culture flasks (VWR, UK). Cells were maintained in AECGM until confluent. Passage number was less than 2.

### Angiotensin-converting enzyme 2

Master stock PM_10_, and CSE (100%) were thawed, thoroughly vortexed and suspended (final PM_10_ concentration 1 to 20 μg/mL, and CSE 5%) in DPBS (2% FBS) containing 2 × 10^5^ A549/ human primary nasal epithelial cells for 2 h at 37°C. Medium controls were incubated with the same volume of DPBS (2% FBS), without PM_10_ or 5% CSE. Cells were washed twice and stained with either anti-ACE2 (Abcam, UK – ab189168, ab272690) or isotype control primary antibodies (Abcam, Ab171870), for 1 h at room temperature. The epithelial cell marker E-cadherin (Abcam, Ab1416) was included in all reactions. Cells were then washed and stained with secondary antibodies conjugated to Alex Fluor 488 for ACE2/isotype expression (Abcam, Ab150077), or APC for E-cadherin expression (Abcam, Ab130786), for 30 min at room temperature in the dark. Cells were finally washed, and ACE2 expression measured using the BD FACS canto II flow cytometer (BD Biosciences, California US). E-cadherin positive cells were selected to exclude cell debris and the median fluorescent intensity (MFI) of fluorescein isothiocyanate (FITC) calculated, adjusting for the isotypic control.

### Statistical analysis

Data are summarised as median (IQR), and analysed by Kruskal-Wallis test with Dunn’s multiple comparisons test. Data are from at least 4 separate experiments.

Analysis were performed using Prism 8 (GraphPad Software Inc., La Jolla, CA, USA) and P<0.05 was considered statistically significant.

## Results

Culture of A549 cells with fossil-fuel derived PM_10_ (0 to 20 μg/mL) for 2 h resulted in a concentration-dependent increase in ACE2 expression, with significant increase at both 10 μg/mL and 20 μg/mL (n=5, P<0.05, P<0.01 vs. medium control, Figure 1). At 20 μg/mL ACE2 increased by 16150 fold (IQR 2577 to 64758).

**Figure 1.**
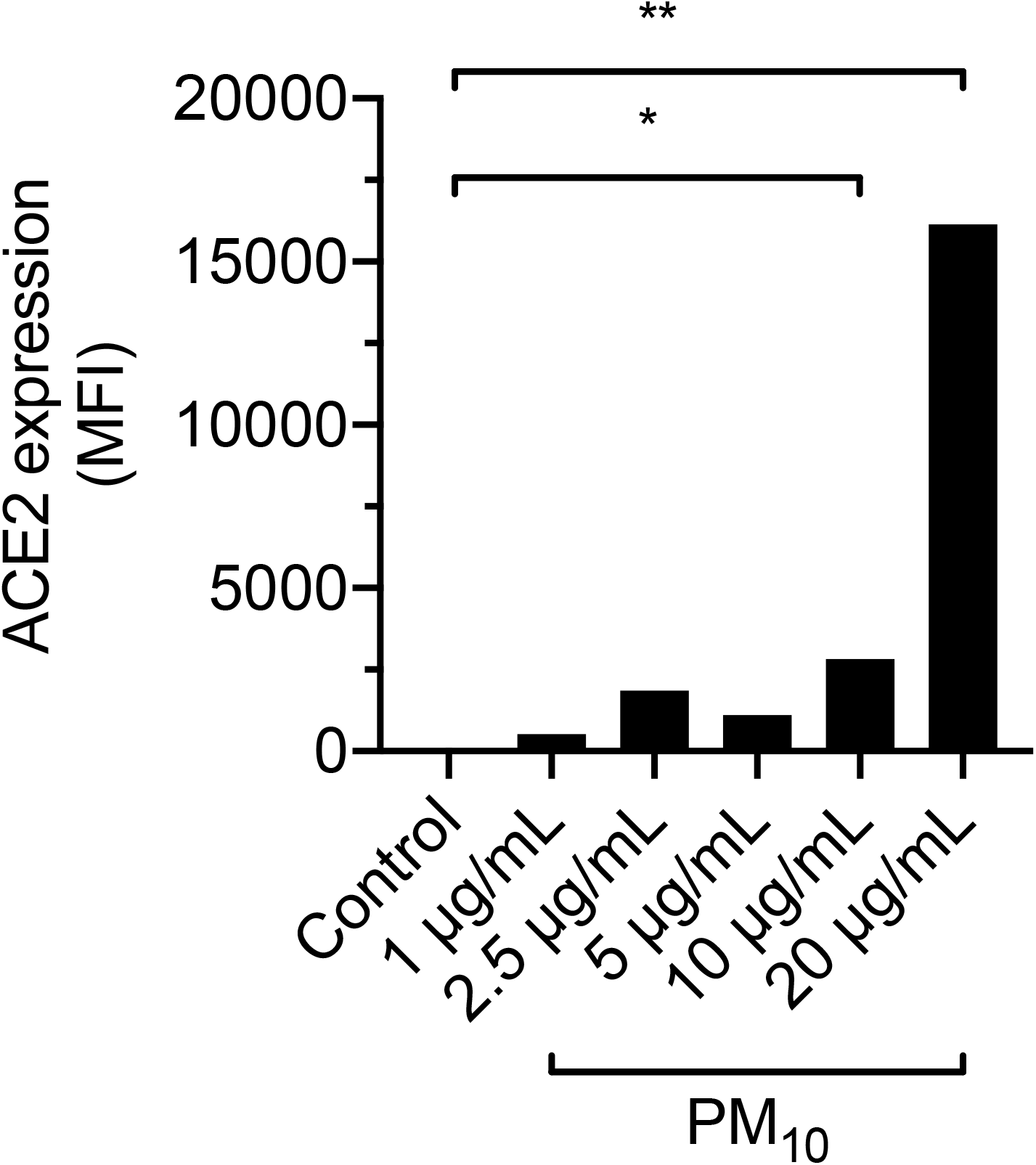
Effect of traffic-derived particulate matter less then 10 microns in aerodynamic diameter (PM_10_) from Marylebone Road, London (UK) on angiotensin-converting enzyme 2 (ACE2) expression in A549 cells. Cells were cultured with PM_10_ for 2 h. ACE2 expression is expressed as median fluorescent intensity (MFI) adjusted for isotypic antibody control. Incubation of cells with PM_10_ at 10 μg/mL and 20 μg/mL increases ACE2 expression (*P <0.05, **P<0.01 vs. medium control). Column represent median from 5 separate experiments. Data are compared by Kruskal-Wallis test and Dunn’s multiple comparisons test.

Using a single concentration of PM_10_ of 10 μg/mL, ACE2 expression increased in A549 cells (MFI; control vs. PM_10_, n=6, 470 (0.1 to 1114) vs. 6216 (5071 to 8506), P<0.01, Figure 2A). PM_10_ also increased ACE2 expression in human primary nasal epithelial cells (MFI, n=4, 0 (0 to 591) vs. 4000 (2610 to 7853), P<0.05, Figure 2B).

**Figure 2.**
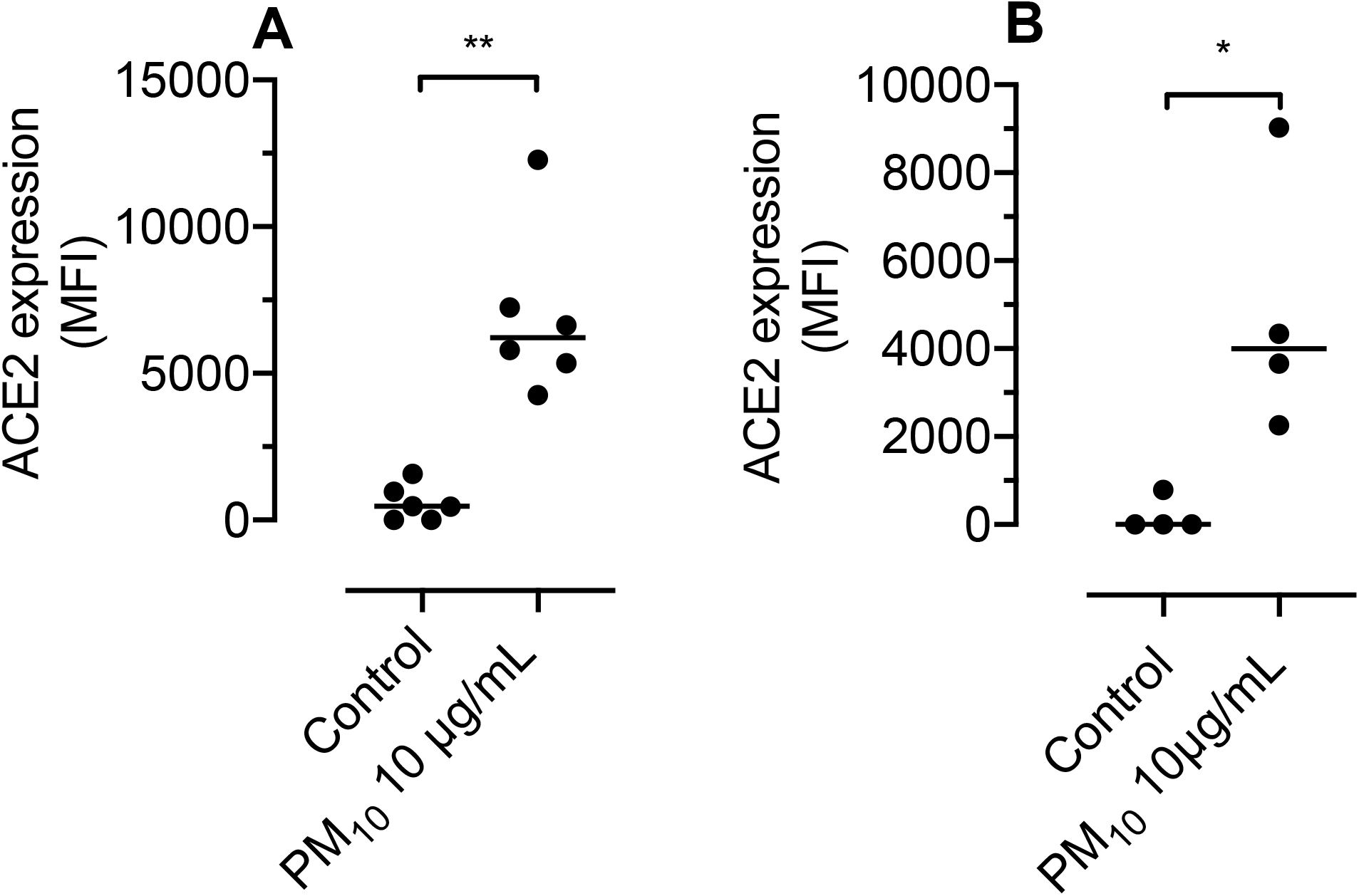
Angiotensin-converting enzyme 2 (ACE2) expression after culture of cells with 10 μg/mL PM_10_ for 2h; A) confirming increased expression in A549 cells. *P<0.01 vs. medium control, and B) increased expression in human primary nasal epithelial cells. *P< 0.05 vs. medium control. Bar represent median. Data are from 4 to 6 separate experiments, and are compared by Mann-Whitney test.

Culture of A549 cells with 5% CSE, a putative positive control, increased ACE2 expression (MFI, n=4, 0 (0 to 28) vs. 9088 (7557 to 15831), P<0.05, Figure 3).

**Figure 3.**
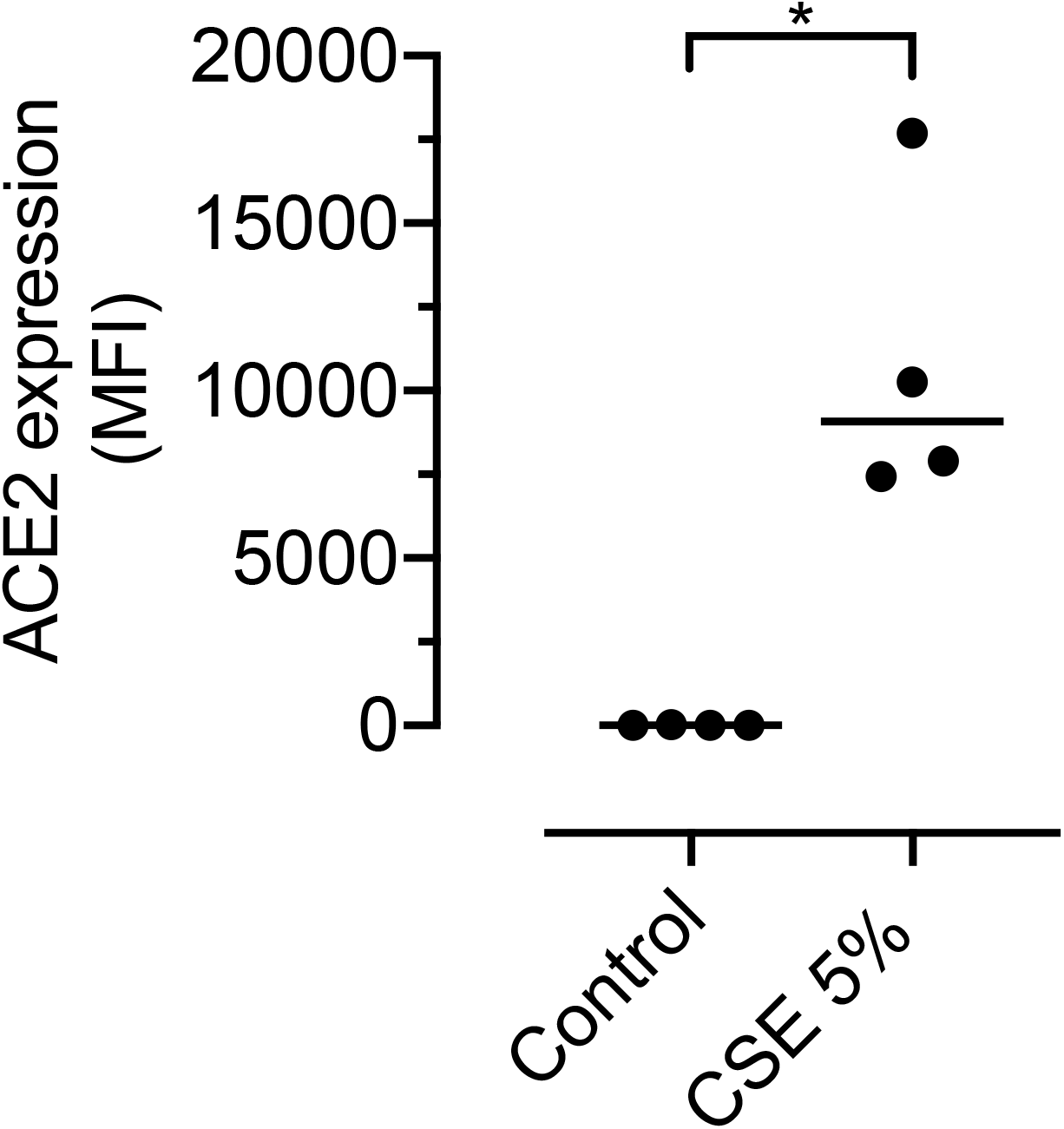
Angiotensin-converting enzyme 2 (ACE2) expression after culture of A549 cells with 5% cigarette smoke extract for 2 h. *P<0.05 vs. medium control. Bar represent median. Data are from 4 separate experiments, and are compared by Mann-Whitney test.

## Discussion

In this study we found that PM_10_, collected next to a major London road dominated by diesel traffic (8), upregulates ACE2 expression in a human type II pneumocyte cell line (A549 cells). We also found that traffic-derived PM_10_ upregulates ACE2 expression in human primary nasal epithelial cells, suggesting that this response occurs throughout the respiratory tract. One strength of the present study is that collection of traffic-derived PM_10_ by a high-volume cyclone obviated the need to extract PM from filters in solution, and we could therefore accurately determine PM_10_ concentrations used in cell culture studies.

Although the effect of PM_10_ on ACE2 expression in human airway cells has not previously been reported, our findings are compatible with an animal study that reported lung ACE2 protein expression in wild type mice increased by 1.3 fold at 2 days post intratracheal instillation of urban PM_2.5_ (11). A putative protective effect of increased pulmonary ACE2 was suggested in this mouse model by complete recovery of PM-induced acute lung injury in wild type mice, and incomplete recovery in ACE2 knockout mice (11). We therefore speculate that increased ACE2 expression may, on one hand, be a beneficial response to PM exposure, but on the other hand presents a Trojan horse to the SARS-CoV-2 virus.

We included CSE as a putative positive control, since Leung *et al* (12) found increased ACE2 gene expression in lower airway brushing from active smokers. Our finding that culture of A549 cells with 5% CSE increases ACE2 expression clearly supports these *in vivo* data, and suggests that cell cultures are a valid method for screening inhaled toxins for capacity to upregulate ACE2 expression. Future screening should include other inhaled toxins including those (e.g. electronic cigarette vapour and welding fumes) previously reported to upregulate expression of platelet activating factor receptor (PAFR) – the host receptor used by *S. pneumoniae* to adhere to airway cells (13)(10).

There are limitations to this study. First, we did not determine whether increased ACE2 expression increases infection of airway cells with SARS-CoV-2. However, evidence for this is provided by reports of; i) an association between smoking and severity to COVID-19 (14) (15), and ii) increased expression of airway epithelial ACE2 in current smokers compared with never smokers (12). By contrast, Jackson *et al* (9) speculated that lower ACE2 mRNA expression in airway brush samples from children with allergic asthma decreases their susceptibility to SARS-CoV-2 infection. Second, we have not identified how PM_10_ upregulates ACE2 expression on airway cells. We have previously reported that PM-induced oxidative stress increases PAFR expression on primary bronchial epithelial cells (16), but the role of oxidative stress on ACE2 expression is as yet unknown. Finally, although the concentration of PM_10_ used in the present study is lower than that used in our previous *in vitro* study of PM and pneumococcal infection of airway cells (16), it remains unclear to what extent 10 μg/mL reflects *in vivo* exposure. Assessment of ACE2 expression in nasal brushings from individuals changing from low to high pollution exposure, for example during and after the COVID-19 lockdown, offers a way of non-invasively validating results from *in vitro* studies.

In conclusion, this study provides the first mechanistic evidence that traffic-derived air pollution increases ACE2 expression in human airway cells and therefore vulnerability to SARS-CoV-2 infection. We conclude that there is biological plausibility for epidemiological studies reporting an association between either PM_10_ or active smoking and COVID-19 disease.

